# Epiphany: predicting Hi-C contact maps from 1D epigenomic signals

**DOI:** 10.1101/2021.12.02.470663

**Authors:** Rui Yang, Arnav Das, Vianne R. Gao, Alireza Karbalayghareh, William S. Noble, Jeffrey A. Bilmes, Christina S. Leslie

**Affiliations:** Memorial Sloan Kettering Cancer Center, New York, NY, USA; University of Washington, Seattle, WA, USA

## Abstract

Recent deep learning models that predict the Hi-C contact map from DNA sequence achieve promising accuracy but cannot generalize to new cell types and indeed do not capture cell-type-specific differences among training cell types. We propose Epiphany, a neural network to predict cell-type-specific Hi-C contact maps from five epigenomic tracks that are already available in hundreds of cell types and tissues: DNase I hypersensitive sites and ChIP-seq for CTCF, H3K27ac, H3K27me3, and H3K4me3. Epiphany uses 1D convolutional layers to learn local representations from the input tracks, a bidirectional long short-term memory (Bi-LSTM) layers to capture long term dependencies along the epigenome, as well as a generative adversarial network (GAN) architecture to encourage contact map realism. To improve the usability of predicted contact matrices, we trained and evaluated models using multiple normalization and matrix balancing techniques including KR, ICE, and HiC-DC+ Z-score and observed-over-expected count ratio. Epiphany is trained with a combination of MSE and adversarial (i.a., a GAN) loss to enhance its ability to produce realistic Hi-C contact maps for downstream analysis. Epiphany shows robust performance and generalization to held-out chromosomes within and across cell types and species, and its predicted contact matrices yield accurate TAD and significant interaction calls. At inference time, Epiphany can be used to study the contribution of specific epigenomic peaks to 3D architecture and to predict the structural changes caused by perturbations of epigenomic signals.

## Introduction

In vertebrate genomes, the three-dimensional (3D) hierarchical folding of chromatin in the nucleus plays a critical role in the regulation of gene expression, replication timing, and cellular differentiation [1, 2]. This 3D chromatin architecture has been elucidated through genome-wide chromosome conformation capture (3C) assays such as Hi-C, Micro-C, HiChIP, and ChIA-PET [3, 4, 5, 6] followed by next generation sequencing, yielding a contact matrix representation of pairwise chromatin interactions. Early Hi-C analyses revealed an organization of ∽ 1Mb self-interacting topologically associating domains (TADs) that may insulate within-TAD genes from enhancers outside of TAD boundaries [7]. High-resolution 3C-based studies have mapped regulatory interactions, often falling within TADs, that connect regulatory elements to target gene promoters [8, 9].

Over the past decade, large consortium projects as well as individual labs have extensively used 1D epigenomic assays to map regulatory elements and chromatin states across numerous human and mouse cell types. These include methods to identify chromatin accessible regions (DNase I hypersensitive site mapping, ATAC-seq) as well as transcription factor occupancy and histone modifications (ChIP-seq, CUT&RUN). While at least some of these 1D assays have become routine, mapping 3D interactions with Hi-C remains relatively difficult and prohibitively costly, and high-resolution contact maps (5 kb resolution, 2 billion read pairs) are still only available for a small number of cell types. This raises the question of whether it is possible to train a model to accurately predict the Hi-C contact matrix from more easily obtained 1D epigenomic data in a cell-type-specific fashion. Such a model could ultimately be used to predict how perturbations in the 1D epigenome—including deletion of TAD boundaries or inactivation of distal regulatory elements—would impact 3D organization.

Initial machine learning methods to predict Hi-C interactions from 1D epigenomic data or DNA sequence took a pairwise approach, treating each interacting or non-interacting pair of genomic bins as an independent training example [10, 11, 12]. For example, HiC-Reg [10] used a random forest regression model to predict the Hi-C contact signal from epigenomic features of the pair of anchoring genomic intervals. Two more recent models, DeepC [13] and Akita [14], respectively predict ‘stripes’ or submatrices of the Hi-C contact matrix from DNA sequence, capturing the non-independence of interaction bins. Neither method uses epigenomic data as an input signal. DeepC [13] presented a transfer learning framework by pre-training a model to predict epigenomic marks from DNA sequence in order to learn useful local sequence representations, then fine-tuning the model to predict the Hi-C contact map. Akita [14] designed a deep convolutional neural network to predict the Hi-C contact maps of multiple cell types from DNA sequence. These prior studies represent a significant advance in predicting 3D genomic structure, and the DeepC and Akita models demonstrated some success in predicting the impact of sequence perturbations like structural genetic variants on local chromatin folding. However, there are also clear limitations to these approaches. Models that start with DNA sequence need considerable computational resources to extract and propagate useful information from base-pair resolution to megabase scale. More importantly, by learning mappings from only DNA sequence to Hi-C contact map data in the training cell types—and therefore lacking any cell-type-specific feature inputs—the resulting models cannot generalize to new cell types that are not seen in training. In fact, it has also been observed that sequence-based models capture very limited cell-type-specific information about 3D genomic architecture even across the training data and instead predict similar structures in every cell type [14].

Here we propose a novel neural network model called Epiphany to predict the cell-type-specific Hi-C contact map from five commonly generated epigenomic tracks that are already available for a wide number of cell types and tissues: DNase I hypersensitive sites and CTCF, H3K27ac, H3K27me3, and H3K4me3 ChIP-seq. Epiphany uses 1D convolutional layers to learn local representations from the input tracks as well as bidirectional long short term memory (Bi-LSTM) layers to capture long term dependencies along the epigenome and a generative adversarial network (GAN) architecture to encourage realism. One goal of our study is to predict contact maps that are usable for downstream computational analyses such as TAD and interaction calls. To this end, we assessed model performance using multiple normalization and matrix balancing techniques including Knight-Ruiz (KR) [15], iterative correction (ICE) [16], and HiC-DC+ [17] Z-score and observed-over-expected count ratio. Epiphany is trained with a combination of mean-squared error (MSE) and adversarial loss to enhance its ability to produce realistic Hi-C contact maps for downstream analysis. The adversarial loss is calculated using a simultaneously trained GAN-style discriminator network, which distinguishes real contact maps from predicted ones, and helps the model to improve its prediction quality. Epiphany shows robust performance and generalization abilities to held-out chromosomes within and across cell types and species, and its predicted contact matrices yield accurate TAD and significant interaction calls. At inference time, Epiphany can be used to study the contribution of specific epigenomic signals to 3D architecture and to predict the structural changes caused by perturbations of epigenomic signals.

## Results

### Epiphany: A CNN-LSTM trained with an adversarial loss accurately predicts Hi-C contact maps

Epiphany uses epigenomic signals (DNaseI, CTCF, H3K27ac, H3K27me3, H3K4me3) to predict normalized Hi-C contact maps. Epigenomic signals are extracted at 100bp resolution from normalized .bigWig files without applying a peak calling step. Hi-C contact maps were initially binned at 10 kb resolution and normalized using the HiC-DC+ package [17] to produce Z-scores and observed-over-expected count ratios, Juicer Tools [18] for KR normalization, and HiCExplorer [19] for ICE normalization. The normalization approaches provided by HiC-DC+ are estimated from a negative binomial regression that is estimated directly from count data and adjusts for genomic distance and other covariates.

Epiphany consists of two parts: a generator to extract information and make predictions, and a discriminator to introduce adversarial loss into the training process (**Fig. 1A** and **Methods**). In the generator, we first used a series of convolution modules to featurize epigenomic information in a sliding window fashion. For one output vector, which covers a distance of 1Mb orthogonal to the diagonal, we used a window size of 1.4 Mb centered at the corresponding region as input (**Fig. 1B**). Then a Bi-LSTM layer was employed to capture the dependencies between output vectors, so that a total of 3.4 Mb input were processed in one pass for prediction of 200 output vectors. At the end, a fully connected layer was used to integrate signals and make the final prediction. We also introduced an adversarial loss and a discriminator, which consists of several convolution modules that are applied during training and pushes the generator to produce realistic samples (**Fig. 1C**).

**Figure 1:**
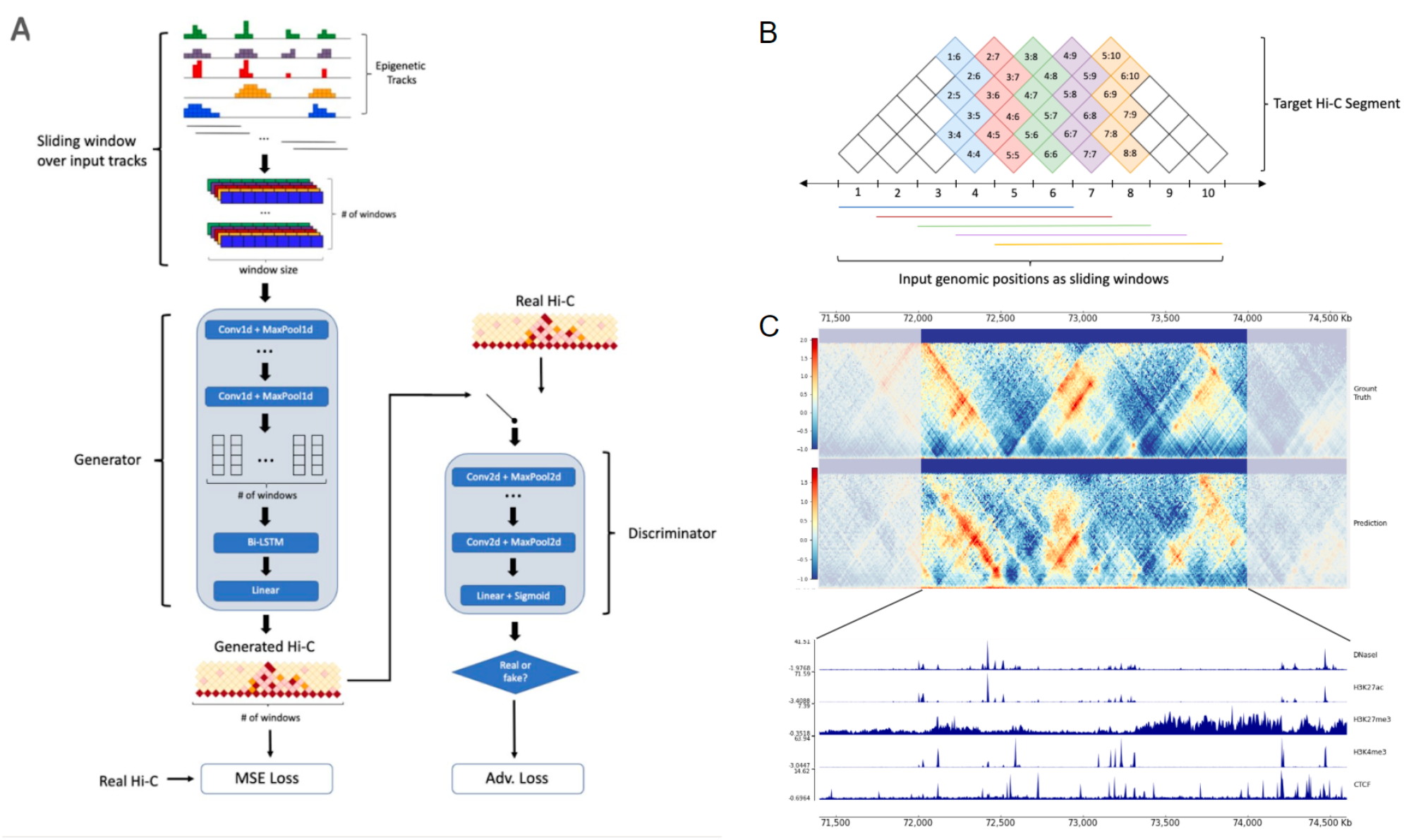
Epiphany employs long short-term memory and adversarial loss to predict the Hi-C contact map. **(A)** Architecture of Epiphany. Epigenomic signal track are first presented to the model in a sliding window fashion, with window size of 1.4 Mb and step size of 10 kb. During training, we take a total length of 3.4 Mb of the input (200 windows) in one pass. In the generator, the processed input data are first featurized by convolution modules, followed by a Bi-LSTM layer to capture the dependencies between nearby bins. After a fully connected layer, the predicted contact map is generated. An MSE loss between the predicted map and the ground truth is calculated in order to train the generator to predict correct structures. To mitigate the overly-smoothed predictions by the pixel-wise losses, we further introduced a discriminator and adversarial loss. The discriminator consists of several convolution modules, and an adversarial loss was calculated to enable the model to generate highly realistic samples. We trained Epiphany with a combined loss of these two components. **(B)** An illustration of prediction scheme. The first window of input data (blue horizontal line, 1.4 Mb) is used to predict a vector on the Hi-C contact map that is orthogonal to the diagonal (blue bin vector, covers 1Mb from the diagonal). During training, a 3.4 Mb length of input are processed using sliding windows (200 windows) in one pass, and 200 consecutive vectors are being predicted. **(C)** An example region of input epigenomic tracks (bottom), target Hi-C map (top row), and predicted Hi-C map (second row).

Given the sequential nature of Hi-C contact maps, interactions on consecutive output vectors are unlikely to be independent from one another. We found that Bi-LSTM layers introduce strong dependencies between the output vectors, which allows Epiphany to leverage structures that span multiple genomic positions in Hi-C maps (such as edges of TADs). Furthermore, Bi-LSTM layers overcome the limitation of convolutional neural networks (CNNs) by enabling each output vector to make use of important signals beyond the input window. This is conducive to studying the contribution of distal regulatory elements towards 3D genome structures and reduces the sensitivity of model performance to the choice of window size.

Past approaches that predict the 3D genome structure from 1D inputs use pixel-wise MSE to quantify the similarity between predicted and ground truth Hi-C maps. However, pixel-wise losses for images have been shown by the computer vision community to be overly sensitive to noise [20] and to yield blurry results when used as objectives for image synthesis [21, 22]. In the context of predicting Hi-C maps, MSE loss can over-penalize poor performance on featureless, noisy regions while failing to penalize underestimation of significant interactions. These issues can be mitigated with an adversarial loss, which enables the model to generate highly realistic samples while circumventing the need to explicitly define similarity metrics for complex modalities of data. Thus, Epiphany is trained using a convex combination of MSE loss and adversarial loss. A parameter *λ* was introduced to balance the proportion of MSE loss and adversarial loss, and the loss function was defined as

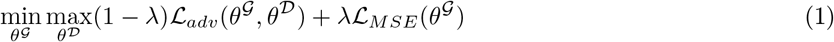

where *λ*ℒ_*adv*_(*θ*^*𝒢*^, *θ*^*𝒟*^) is the adversarial loss, and ℒ_*MSE*_(*θ*^*𝒢*^) is the MSE between the predicted contact map and ground truth. Intuitively, the MSE loss ensures that the Hi-C maps predicted by Epiphany are aligned with their corresponding epigenomic tracks, while the adversarial loss ensures that the predictions are realistic. We find that using this customized training objective yields Hi-C maps that can be directly processed by commonly used downstream analysis tools.

### Epiphany accurately predicts the Hi-C contact map

We first benchmarked the model at 10 kb resolution to compare between two loss functions: MSE only and the convex combination of MSE and adversarial loss. Both losses use the observed-over-expected count ratio normalization based on HiC-DC+. Models were trained on data from the GM12878 ENCODE cell line, with chr3, 11, and 17 as completely held-out chromosomes. Epiphany demonstrates good performance for both the Pearson and Spearman correlation metrics using the observed-over-expected count ratio (**Table 1**), while MSE produced higher correlations than the convex combination of MSE and adversarial loss. However, we observed that the high correlations from MSE trained models were associated with blurriness in the predicted contact maps (**Fig. 2B**), whereas the correlations produced by the combined loss models may have been slightly diminished due to small deviations in the sharper predictions. We also found that downstream algorithms such as TAD or significant loop callers would not function properly on such blurry maps. Therefore, we reasoned that correlation may not be an appropriate evaluation metric and decided to use the combined loss (MSE+adversarial loss) for downstream analysis.

**Table 1:** Mean Pearson and Spearman correlation for different normalization methods

| Normalization Method | $\lambda$ | Pearson (all) | Pearson (train) | Pearson (test) | Spearman (all) | Spearman (train) | Spearman (test) |
| --- | --- | --- | --- | --- | --- | --- | --- |
| **Obs/Exp** | 0.95 | 0.7408 | 0.7687 | 0.5636 | 0.6899 | 0.7191 | 0.5048 |
| Obs/Exp | 1 (MSE only) | 0.7833 | 0.8045 | 0.6494 | 0.7381 | 0.7605 | 0.5963 |
| Z-score | 0.95 | 0.6881 | 0.7222 | 0.4722 | 0.6695 | 0.7034 | 0.4544 |
| KR | 0.35 | 0.7289 | 0.7510 | 0.5889 | 0.5909 | 0.6135 | 0.4477 |
| ICE | 0.35 | 0.8108 | 0.8288 | 0.7028 | 0.6631 | 0.6852 | 0.5303 |

**Figure 2:**
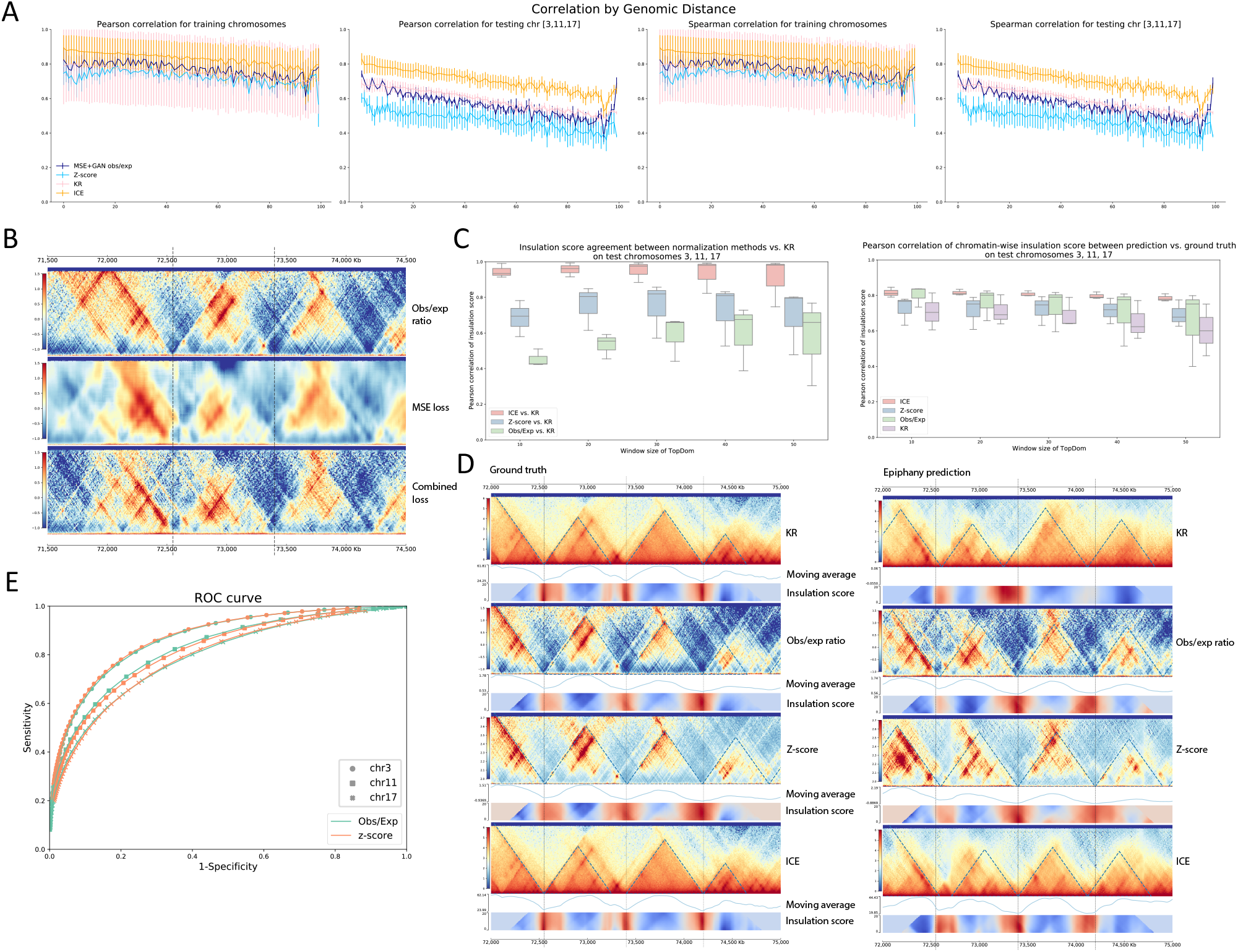
Epiphany-predicted contact maps identify TADs and significant interactions. **(A)** Epiphany performance using correlation by genomic distance for different normalization approaches. From left to right: Pearson correlation on training chromosomes, on testing chromosomes (chr3, 11, 17), and Spearman correlation on training and on testing chromosomes. Dark blue shows the performance of HiC-DC+ observed-over-expected count ratio, light blue shows HiC-DC+ Z-score, pink shows KR normalization, and yellow shows ICE normalization. **(B)** A visual comparison of the ground truth contact map (chr17:70,670,000-73,880,000, top row), blurry prediction made by MSE trained model (middle row), and more realistic prediction by combined loss (bottom row). **(C)** Top: Agreement of insulation score between different normalization methods vs. KR normalization on test chromosomes. Insulation scores were calculated using TopDom with different window sizes (X-axis) on ground truth contact maps with different normalization methods. KR normalization was used as the gold standard, and a Pearson correlation (Y-axis) was calculated to measure the agreement between each normalization method vs. KR (red: ICE vs. KR, blue: HiC-DC+ Z-score vs. KR, green: HiC-DC+ obs/exp vs. KR). Bottom: Pearson correlation of insulation score between predicted contact map vs. corresponding ground truth of the same normalization (red: ICE, blue: HiC-DC+ Z-score, green: HiC-DC+ obs/exp, purple: KR). **(D)** Left: Ground truth contact maps of different normalization methods (from top to bottom: ICE, HiC-DC+ obs/exp, HiC-DC+ Z-score, KR). Blue dashed lines denotes the TAD calls with window size of 50 on each contact map, and black dashed lines are the TAD boundaries called from KR normalized contact map. Right: Predicted contact maps of different normalization. **(E)** ROC curve of significant interactions between prediction contact maps vs. ground truth for the three test chromosomes for HiC-DC+ obs/exp ratios (green) and for Z-scores (orange).

We then tested the robustness of Epiphany with various normalization methods, including KR normalization, ICE normalization, and Z-scores from HiC-DC+. All models were set up with the same training approach as before, where chr3, 11, and 17 were used as held-out chromosomes and models were trained with the combined loss. Epiphany shows robust performance in all normalization methods (**Fig. 2A**). ICE normalization obtained the highest correlations, with an average Pearson correlation of 0.7028 and Spearman correlation of 0.5303 on completely held-out chromosomes (**Table 1**).

To explore the capacity of Epiphany to capture key structures in genome architecture, we next evaluated the ability of Epiphany predictions to recover TAD boundaries. For all normalization methods and their predictions, we called TAD boundaries using TopDom [23] with window sizes ranging from 10 to 50 (corresponding to 100 kb to 500 kb regions). Because TAD calls depend on the normalization method, we first used KR normalization as the gold standard and compared TAD insulation scores computed on ground truth data on the test chromosomes using different normalization methods. Note that we chose to compare insulation scores rather than TAD boundaries, since the latter relies on finding local extrema in the insulation score signal and therefore can be unstable. Among all these methods, ICE had the highest consistency with KR, followed by Z-scores calculated from HiC-DC+. The observed-over-expected count ratios had the least consistency and showed large variation over the three test chromosomes (**Fig. 2C**, left). We then compared the insulation score calculated from the Epiphany-predicted contact maps trained with different normalization methods vs. the corresponding ground truth on the test chromosomes. ICE showed robust predictions on all test chromosomes, whereas HiC-DC+ observed-over-expected count ratio normalization displayed strong mean performance but had larger variance, especially for larger window sizes. HiC-DC+ Z-scores and KR normalization showed lower consistency between predicted vs. ground truth insulation scores (**Fig. 2C**, right). From a visual comparison of ground truth and predicted contact maps with different normalization approaches, we could see Epiphany consistently predicts accurate TAD structures (**Fig. 2D**). Overall, this analysis suggests that, for accurate prediction of TAD structure, Epiphany trained on ICE normalized contact maps gave the best performance, with HiC-DC+ observed-over-expected count ratio as runner-up.

One advantage of HiC-DC+ normalization is that it readily allows the comparison of significant interactions between predicted contact maps and ground truth. HiC-DC+ [17] fits a negative binomial regression using genomic distance, GC content, mappability and effective bin size based on restriction enzyme sites to estimate the expected read count for each interaction bin, which allows an assessment of significance of the observed count. For convenience, we defined the significant interactions as ground truth Z-scores greater than 2. Significant interactions were called with various thresholds from test chromosomes on predicted contact maps using Z-scores and observed-over-expected count ratio, yielding the ROC curves for each test chromosome (**Fig. 2E**). The average AUC is 0.7639 for the two models, suggesting solid performance at a difficult task.

### Epiphany shows robust performance at finer resolution

Due to good overall performance and the ability to directly identify significant interactions, we chose observed-over-expected count ratios rather than Z-score from HiC-DC+ for further analysis. We again trained Epiphany to predict interactions within 1Mb from the diagonal at 5 kb resolution. Epiphany showed robust performance at 5 kb resolution, with an average Pearson correlation of 0.5625 and Spearman correlation of 0.5270. In addition to the distance-dependent correlations, we also used both MSE loss and insulation scores calculated from HiCExplorer [19] to evaluate model performance. Since Epiphany jointly predicts multiple interaction vectors, the model can predict a submatrix of the contact map that covers a 2Mb distance along the diagonal (400 vectors for 5 kb resolution) and up to 1Mb from the diagonal. We calculated the average MSE loss between the predicted submatrix vs. ground truth as well as Pearson correlation between insulation scores calculated from the corresponding submatrices. Results for all 2Mb submatrices from the three held-out chromosomes (chr3, 11, 17, **Fig. 3B**) show that Epiphany displays consistent prediction performance across held-out chromosomes with diverse length and gene densities. In particular, 84.4% (173 out of 203) of submatrices have insulation correlation higher than 0.50. Epiphany showed robust performance in most regions along the genome but sometimes produced inaccurate predictions at regions without clear signals or in low mappability regions. (**Fig. 3A**).

**Figure 3:**
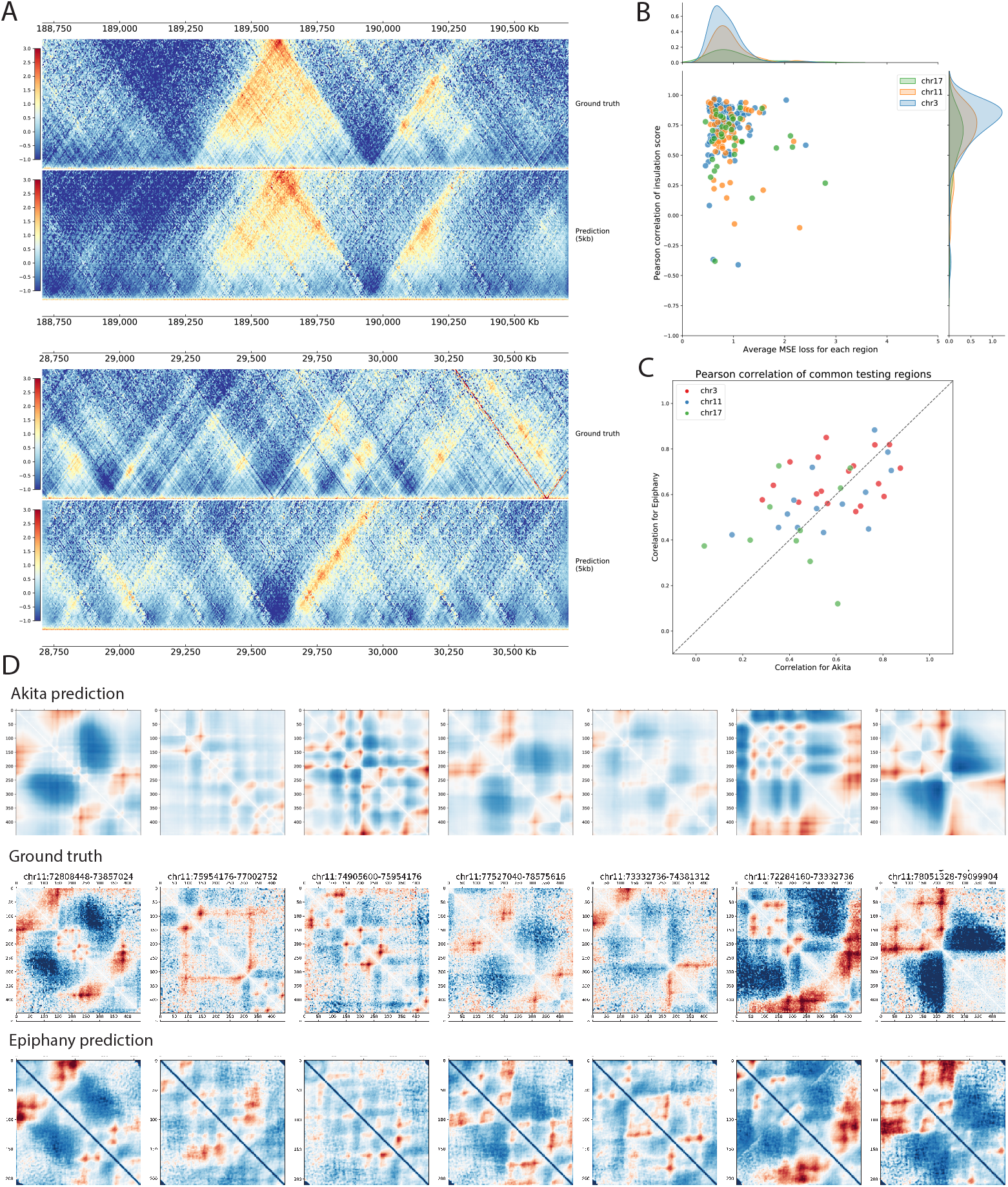
Epiphany achieves state-of-the-art performance at fine resolution. **(A)** Epiphany performance evaluation at 5 kb resolution. Top: one of the best predicted submatrices (chr3:188,610,000-190,610,000) with ground truth matrix on the top, and predicted matrix on the bottom. Bottom: one of the problematic matrices (chr17:28,705,000-30,705,000) predicted by Epiphany. **(B)** Evaluation of predicted submatrices. X-axis denotes the average MSE loss between predicted matrix and ground truth, and Y-axis shows the Pearson correlation of insulation score of the 2 Mb region. Dots are colored by chromosomes, and density plots for dot distribution are added on the side. **(C)** Model performance comparison between Epiphany and Akita on 42 common regions between Akita held-out test regions and our test chromosomes (chr3, 11, 17). X-axis shows the Pearson correlation of Akita prediction vs. ground truth, and Y-axis shows the correlation of Epiphany. Epiphany was re-trained using data with the same normalization steps of Akita at 5 kb resolution, and Akita predictions were average-pooled into 4096 bp resolution for better comparison. Dots are colored by chromosomes. **(D)** Visual comparison of Akita prediction (2048 bp resolution, top row), ground truth matrices (2048 bp resolution, middle row), and Epiphany prediction (5 kb, bottom row).

We also compared Epiphany with Akita [14] on common test regions, restricting our evaluation to regions that were held out by Akita and overlap with our test chromosomes. We binned the Hi-C contact map at 5 kb resolution and followed the normalization approaches suggested in the Akita study (Methods). Epiphany was re-trained using the training chromosomes as before (all chromosomes except for chr3, 11, 17) and evaluated on the 42 test regions from Akita’s held-out set falling in our test chromosomes. Akita’s predictions at 2048 bp resolution were average-pooled to 4096 bp in order to obtain relatively consistent resolution. For each test region, we calculated the Pearson correlation between predicted contact matrices and ground truth for both Akita and Epiphany (**Fig. 3C**). We also visually compared the predictions of Akita and Epiphany with ground truth on the held-out examples (**Fig. 3D**). Quantitatively and qualitatively, both models showed similar performance.

### Epiphany predicts cell-type specific 3D structure

Since Epiphany uses epigenomic marks as input, it can potentially generalize to predict cell-type-specific 3D structures in a new cell type. We first compared Epiphany’s cell-type-specific predictions with those of Akita, where five different cell types were simultaneously predicted in a multi-task framework. We selected H1ESC and GM12878 from these five cell types for the comparison. Akita’s cell-type-specific predictions were directly obtained from its multi-task output. Epiphany was trained on GM12878 and evaluated on H1ESC test chromosomes (chr3, 11, 17) at inference time. We checked the visual comparison of Akita and Epiphany cell-type-specific predictions relative to their respective ground truths and also calculated the absolute difference for ground truth and predictions between the two cell types (**Fig. 4A**). The results suggest that Epiphany, which was trained only on GM12878 data, can generalize to a new cell type and accurately predict the differential structure between cell types based on cell-type-specific 1D epigenomic data. By contrast, the DNA-sequence-based Akita model, although trained on Hi-C/Micro-C data in these and other cell types, largely predicts the same 3D structure in GM12878 and H1ESC.

**Figure 4:**
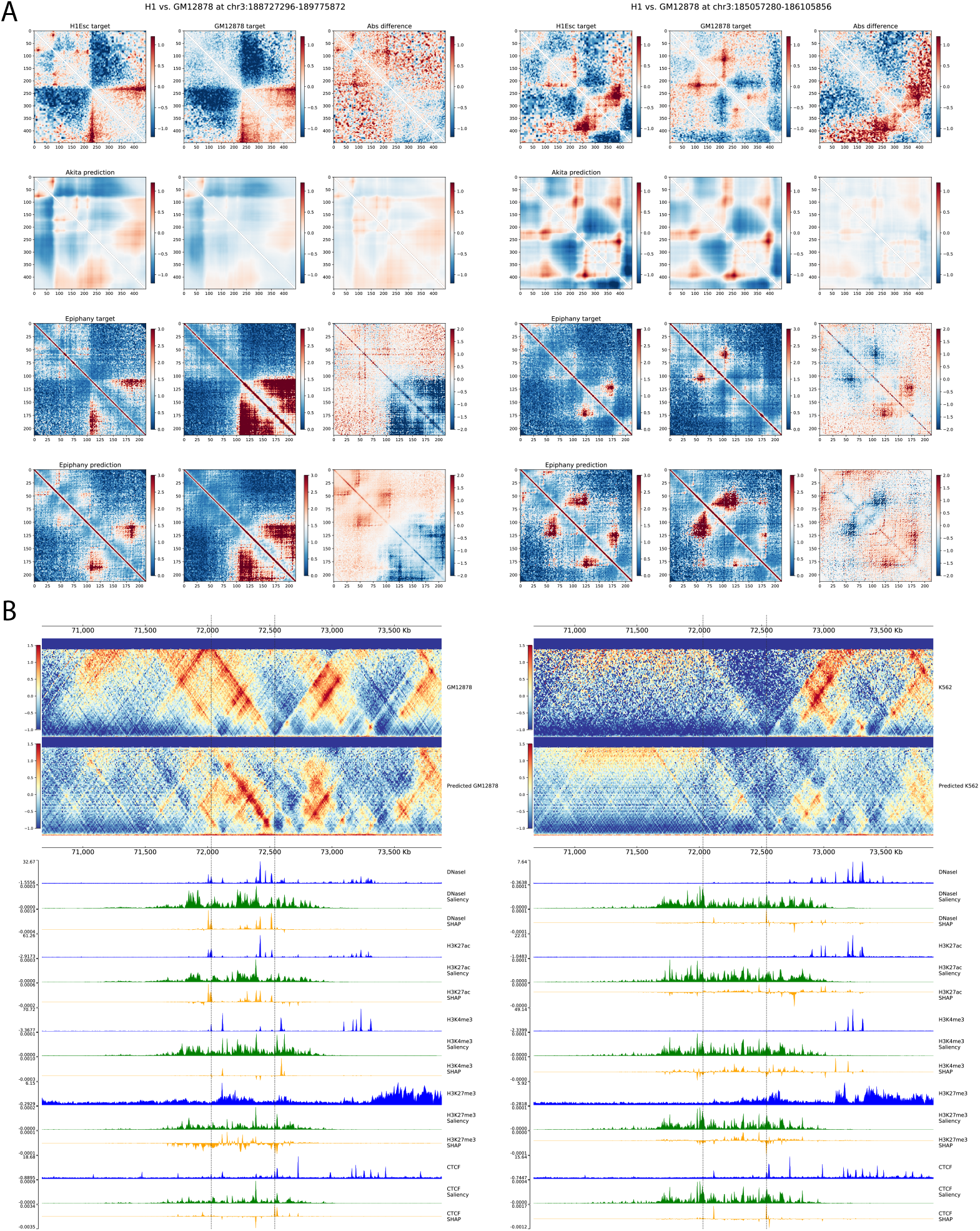
Epiphany accurately predicts cell-type-specific 3D structures. **(A)** Two examples (chr3:188,727,296-189,775,872) and (chr3:185,057,280-186,105,856) of cell-type specific predictions in H1ESc and GM12878. Two regions are selected from the overlapped region of Akita held-out test set and Epiphany’s test chromosomes. Columns from left to right: contact map in H1ESc, same region in GM12878, and the absolute difference between the two cell types (H1ESc-GM12878). Rows from top to bottom: Ground truth matrices with Akita normalization, Akita prediction, ground truth with HiC-DC+ observed-over-expected count ratio, Epiphany prediction of observed-over-expected count ratio. Akita predictions were obtained from the multi-task output, and Epiphany predictions were generated with model trained on GM12878. **(B)** Cell type specific prediction at a differential region between GM12878 and K562. On the left is the ground truth matrix (top) and predicted matrix (middle), followed by epigenomic input tracks (blue), Saliency score (green), and SHAP values (yellow) for feature attributions. On the right is the predictions for K562. Epiphany was trained in GM12878 in training chromosomes, and predicted both cell types for test chromosomes.

We next explored the ability of Epiphany to identify the contribution of cell-type-specific epigenomic input features to differential 3D structures using feature attribution. In recent years, feature attribution has become a powerful tool to study the contribution of input features to prediction of a specific output. For each interaction bin in the predicted contact map, we first calculated the saliency score [24], which is a gradient-based attribution on input values. We then calculated the SHAP value [25] with baseline signals equal to zero, which highlights the contribution of epigenomic peaks to a specific output. We compared a region (chr17:70,500,000-73,500,000) with differential interactions between GM12878 and K562 (**Fig. 4B**). Epigenomic signals between chr17:72,000,000-72,500,000 in GM12878 contributed to the prediction of the highlighted interaction, while the absence of signals in K562 input led to the correct prediction of a weak interaction.

### Ablation analyses suggests redundancies between 1D inputs

In the previous cell-type-specific analysis, distal H3K4me3 peaks gained importance in the K562 prediction when there were no signals at the anchors of the investigated interaction (**Fig. 4B**). We wondered whether features from different epigenomic tracks could compensate for each other in predicting interactions and more generally whether there exist redundancies between the input tracks.

We performed a feature ablation experiment to address these questions. Instead of including all five epigenomic tracks as input, we re-trained the model with one or several of the tracks completely masked as zero. We reasoned that re-training the model rather than masking a specific input region at test time could better serve our goal. For example, using a model trained on all five input tracks, if we simply masked one important peak from DNaseI track during test time, we expected that the model would inevitably fail to predict the corresponding interactions. However, if we re-trained the model with the entire DNaseI track masked, we expected the model to identify alternative signals from other tracks during training and potentially retain the ability to predict these interactions.

Indeed, this idea matched our observations from the ablation analysis. We re-trained Epiphany with a) an additional SMC3 ChIP-seq track, b) CTCF track masked as zero, c) DNaseI track masked as zero, d) only CTCF and H3K27ac tracks as training inputs, and compared their predictions with the results using all input (**Fig. 5A**). We found that by removing DNaseI, the model achieved similar performance as using all input tracks. Models with CTCF masked or using only two tracks (CTCF+H3K27ac) showed weaker performance.

**Figure 5:**
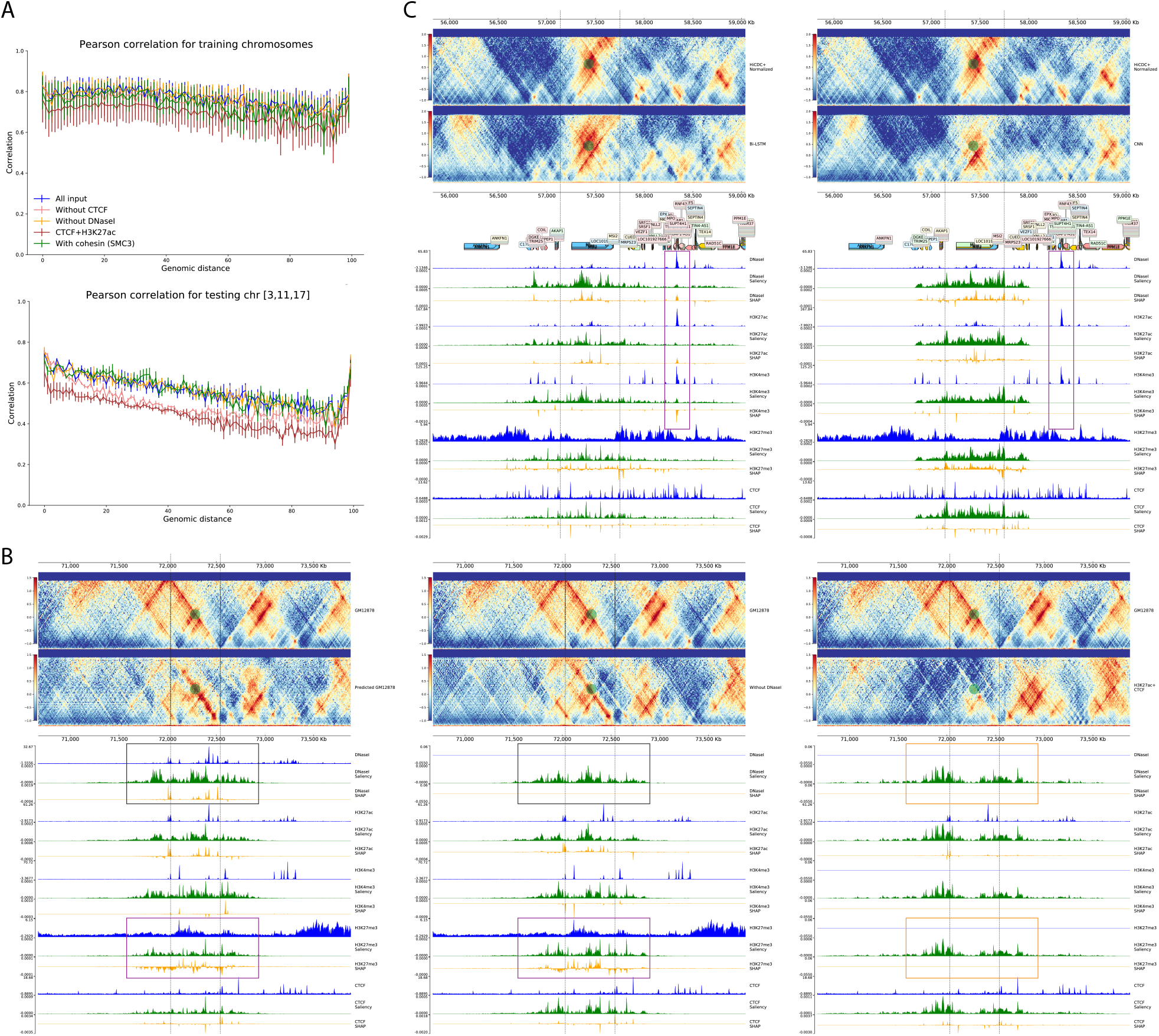
Feature ablation and attribution identify the contribution of epigenomic marks to 3D structure. **(A)** Correlation by distance for feature ablation experiments. Top: Pearson correlation for training chromosomes. Bottom: Pearson correlation for test chromosomes. Blue track for model performance using all 5 epigenomic input tracks; green for training with an additional track SMC3; pink for model trained without CTCF; yellow for without DNaseI, and dark red for model trained with only CTCF and H3K27ac tracks. **(B)** Feature attribution for bin (chr17:72,030,000-72,540,000) with full model (left), model without DNaseI (middle), model with only CTCF and H3K27ac (right). **(C)** Feature attribution for bin (chr17:57,140,000-57,750,000) with Epiphany with Bi-LSTM layer (left) vs. modified Epiphany with convolution layer (right).

As we have seen in previous example (**Fig. 4B**), DNaseI and H3K27ac contributed to the differential predictions between GM12878 and K562 at the region chr17:70,670,000-73,880,000. We therefore compared the prediction for this region using a model trained with all input tracks, without DNaseI, or with CTCF+H3K27ac only (**Fig. 5B**). Epiphany was still able to accurately predict interactions in this region after ablating DNaseI; feature attribution indicated that in place of the DNaseI signal (**Fig. 5B**, grey box), the model gave higher importance to H3K27me3 peaks (purple box) in order to predict the interaction. However, after ablating all signals except for CTCF and H3K27ac, the model failed to find alternative predictive signals and missed the boundary.

### Bi-LSTM layers capture the contribution of distal elements

Given the sequential nature of Hi-C contact maps, interactions on consecutive output vectors are unlikely to be independent from one another. We tested whether the Bi-LSTM layers in Epiphany indeed captured the dependencies between the output vectors better than regular convolutional layers. We made predictions using Epiphany with Bi-LSTM layers and compared with a modified Epiphany where the Bi-LSTM layers were replaced by convolutional layers. We also calculated the saliency score and SHAP values for the bin chr17:57,140,000-57,750,000 (**Fig. 5C**). The interaction at chr17:57,140,000-57,750,000 was better predicted by Epiphany with Bi-LSTM layers, and feature attribution showed that one distal peak at around chr17:58,350,000-58,400,000 contributed to the prediction. Compared with regular convolutional layers, Bi-LSTM layers introduce stronger dependencies between the output vectors and overcome the limitation of CNNs by enabling each output vector to make use of important signals beyond the input window.

### Epiphany predicts perturbations in 3D architecture

Since Epiphany models the contribution of epigenomic signals to 3D structures, we explored whether Epiphany could predict 3D structural changes caused by perturbations to the epigenome. In particular, we considered examples where structural variations eliminate important epigenomic features. Despang *et al*. [26] studied the TAD fusion caused by deletion of CTCF sites in vivo in the mouse embryonic limb bud at the *Sox9-Kcnj2* locus. They used the promoter capture Hi-C data in the E12.5 mouse limb bud to show the structural changes after deleting major CTCF sites (mm9, GSE78109, GSE125294). In WT TAD structures, *Kcnj2* and *Sox9* are separated into two TADs. After deleting four consecutive CTCF sites within a 15 kb boundary region (C1 site mm9 chr11:111,384,818–111,385,832, C2-C4 site chr11:111,393,908-111,399,229), the TAD boundaries disappeared and the two TADs fused together. When all CTCF binding sites between *Kcnj2* and *Sox9* were deleted, they observed a more complete TAD fusion (**Fig. 6A**). These experiments revealed a TAD fusion caused by the deletion of major CTCF sites at the boundaries and within the TAD.

**Figure 6:**
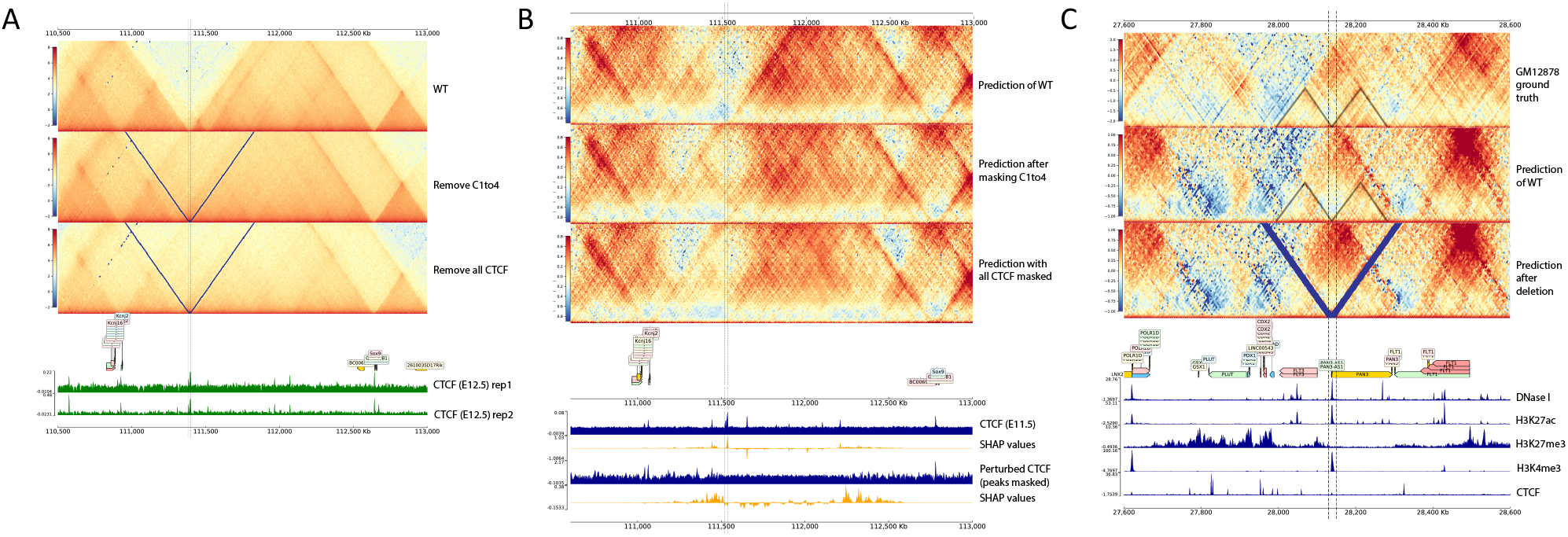
Epiphany predicts TAD boundary changes due to epigenomic perturbations. **(A)** Mouse ES E12.5 Capture Hi-C data from Despang *et al*. [26] for WT (top), 4 CTCF sites depletion (middle), and all CTCF depletion between gene *Kcnj2* and *Sox9* (bottom). Data are publicly available at (GSE78109, GSE125294). Four CTCF sites depleted in the middle figure were at region (C1 site(mm9 chr11:111,384,818–111,385,832), C2-C4 site (chr11:111,393,908-111,399,229), marked with black dashed lines). Data are mapped relative to mm9. **(B)** Epiphany cross-species prediction of structural changes caused by CTCF perturbation. Epiphany was trained using human cell line GM12878, and predicted using mouse limb bud epigenomic data mapped relative to the mm10 assembly. The panel shows Epiphany prediction of WT mES Hi-C map with HiC-DC+ obs/exp ratio normalization (top row), the prediction of TAD fusion after masking CTCF sites at (mm10 chr11:111,520,000-111,540,000) (middle row), and the prediction of further TAD fusion after masking all CTCF peaks between *Kcnj2* and *Sox9* genes (bottom row). Epigenomic tracks at the bottom are showing feature attribution (SHAP value) for highlighted vertical vector in the Hi-C contact map. The upper two tracks are the original CTCF track and corresponding SHAP values. The lower two tracks are CTCF tracks with peaks between *Kcnj2* and *Sox9* masked to the background and the corresponding SHAP values. **(C)** Human GM12878 Hi-C contact map at 5 kb resolution at chr13:27,600,000-28,600,000. Ground truth contact map (top), predicted contact map with unperturbed input (middle), predicted contact map with 20 kb deletion at region chr13:28,130,000-28,150,000 (bottom, deleted region highlighted in dashed line).

We then tested Epiphany’s ability to predict these structural changes after we perturbed the CTCF input track. Epiphany was trained on data from the human cell line GM12878 and used to make cross-species prediction in the mouse embryo. Epigenomic tracks were downloaded from ENCODE [27] and BioSamples [28] for mouse limb tissue aligned to the mm10 assembly. In the WT prediction, Epiphany predicted a strong boundary separating *Kcnj2* and *Sox9* into two TADs. Upon masking the C1-4 CTCF peaks (mm10 chr11:111,520,000-111,540,000) at the boundary and further masking all CTCF sites, Epiphany predicted behavior consistent with the ground truth experiments, where the two TADs gradually merged together (**Fig. 6B**, top). We further explored the relationship between the CTCF peaks and TAD formation using feature attribution methods (**Fig. 6B**, bottom). The SHAP values are calculated and averaged for the bins in the vertical highlighted dashed lines. We can see that CTCF peaks at the boundary contribute to TAD separation in unperturbed prediction, while with the masked CTCF track, the feature attribution scores focus on more distal regions at the boundaries of the fused TAD.

We also evaluated whether Epiphany could predict structural changes caused by genomic deletions. Yang *et al*. [29] observed upregulation of the *FLT3* gene in acute lymphoblastic leukemia (ALL) patients with a 13q12.2 deletion and attributed this gene expression change to chromatin structural reorganization and enhancer hijacking. *FLT3* was found to be controlled by three regulatory elements in the 13q12.2 region: DS1 (chr13:28100363-28100863), the promoter of *FLT3*; DS2 (chr13:28,135,863-28,140,863); and DS3 (chr13:28,268,863-28,269,363) in the intron of *PAN3*. In normal hematopoietic cells, *FLT3* is primarily controlled by the interaction of DS1 and DS2 due to the separation of two TADs (**Fig. 6C**, top, highlighted with dashed lines), where DS2 overlaps with the TAD boundary, and DS3 is located in a nearby TAD. In patients with the 13q12.2 deletion where DS2 was lost, they observed a fusion of the two nearby TADs and a strengthened long-range interaction between DS1 and DS3.

We simulated this deletion by excising the DS2 region from all epigenomic input tracks and predicted the resulting contact matrix with Epiphany. That is, epigenomic signals in all five input tracks for the corresponding region were deleted, and the up- and downstream tracks were concatenated together. Epiphany predicted TAD structures consistent with reported observations. Before 13q12.2 deletion, Epiphany predicted a small TAD separation between *FLT3* and *PAN3* genes at (chr13:27,600,000-28,600,000) region, consistent with the ground truth in GM12878 (**Fig. 6C**, top and middle). After the deletion, Epiphany shows the fusion of the two TADs and increased interactions between the *FLT3* and *PAN3* gene regions (**Fig. 6C**, bottom).

## Discussion

In this study, we developed Epiphany, a neural network model to predict the cell-type specific Hi-C contact map for entire chromosomes up to a fixed genomic distance using commonly generated epigenomic tracks that are already available for diverse cell types and tissues. We showed that Epiphany accurately predicts cell-type-specific 3D genome architecture and shows robust performance for Hi-C different normalization procedures and at different resolutions. Epiphany was able to accurately predict cross-chromosome, cross-cell type and even cross-species 3D genomic structures. From feature ablation and attribution experiments, we showed that Epiphany could be used to interpret the contribution of specific epigenomic signals to local 3D structures. Through in silico perturbations of epigenomic tracks followed by contact map prediction with Epiphany, we were able to accurately predict the cell-type-specific impact of epigenetic alterations and structural variants on TAD organization in previously studied loci.

Although we used five specific epigenomic tracks (DNase I, CTCF, H3K27ac, H3K27me3, H3K4me3) and Hi-C data in this study, we believe that Epiphany could be used as a more general framework to link cell-type-specific epigenomic signals to 3D genomic structures. In the future, we plan to explore different combinations of the epigenomic input tracks to assess their biological and statistical relevance for prediction of 3D structure.

In addition to using the epigenomic information, we also tried to incorporate DNA information into the model. Previous models have used a one-hot encoding of long genomic DNA sequences 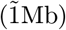, incurring significant computational costs [14, 13]. We therefore tried an alternative strategy of extracting DNA representations from a pre-trained DNABERT model [30], a new method that adapts the state-of-the-art natural language processing model BERT [31] to the setting of genomic DNA. During the pre-training phase, DNA sequences were first truncated to 510 bp length sequences as the ‘sentences’ and further divided into k-mers as the ‘words’ of the vocabulary. The model learns the basic syntax and grammar of DNA sequences by self-supervised training to predict randomly masked k-mers within each sentence. After pre-training, each 510 bp sequence was represented by a 768-length numerical vector. However, since Epiphany covers a 3.4 Mb region as input during training, it was still extremely computationally intensive to directly incorporate the pre-trained representations from DNABERT. We therefore excluded the DNABERT component in order to keep the model relatively light-weighted and concise, although we do not rule out its utility in the future.

Beyond these computational issues, a more conceptual modeling challenge is retaining the ability to generalize to new cell types while also incorporating DNA sequence information. In principle, training on genomic sequence may learn DNA sequence features that are specific to the training cell types and do not generalize to other cell types. Epiphany learns a general model for predicting the Hi-C contact map in a cell type of interest from cell-type-specific 1D epigenomic data, giving state-of-the-art prediction accuracy while allowing generalization across cell types and across species.

## Methods

### Data sources and pre-processing

#### Training and test sets

We used three human cell lines (GM12878, H1ESc, K562) and one mouse cell line (mES) for training and testing the model. All human data (Hi-C, ChIP-seq) were processed using the hg38 assembly and mouse data with mm10. For all experiments, chromosome 3, 11, 17 were used as completely held-out data for testing.

#### Epigenomic data

All input epigenomic tracks including DNaseI, CTCF, H3K4me3, H3K27ac, H3K27me3 for genome assembly hg38 were downloaded from the ENCODE data portal [27]. Data were downloaded as bam files, and the replicates for each epigenomic track were merged using the pysam (https://github.com/pysam-developers/pysam) python module. We then converted merged bam files into bigWig files with deepTools [32] bamCoverage (binSize 10, RPGC normalization, other parameters as default). Genome-wide coverage bigWig tracks were later binned into 100-bp bins, and bin-level signals for the 5 epigenomic tracks were extracted as input data for the model.

#### Hi-C data

High quality and deeply sequenced Hi-C data as .hic format for all human and mouse cell lines were downloaded from 4DN data portal [1]. Data were binned at 5 kb and 10 kb resolution and normalized using multiple approaches. KR normalization was calculated by Juicer tools [18] and ICE normalization by the HiCExplorer package [19]. Observed-over-expected count ratio and Z-score normalizations were calculated by HiC-DC+ [17]. ICE normalization for 5 kb resolution was calculated using Cooler [33], and all additional matrix balancing steps followed the Akita pipeline [14]. For the observed-over-expected count ratios from HiC-DC+, raw counts for interaction bins are modeled using negative binomial regression to estimate a background model, giving an expected count value based on the genomic distance and other covariates associated with the anchor bins (GC content, mappability, effective size due to restriction enzyme site distribution). The observed-over-expected count ratio is then calculated using observed raw counts divided by the expected counts from the HiC-DC+ model.

#### Biological validation data

Capture Hi-C and corresponding CTCF tracks from Despang *et al*. [26] were downloaded from (GSE78109, GSE125294). Data were visualized using Coolbox [34].

### Model and training

#### CNN layers

The input epigenomic tracks were divided into overlapping windows, with a window length of *m* = 14, 000 bins (1.4Mb) and a stride of 1,000 bins (100 kb). We refer to the windowed inputs as *X* = {*x*_1_, …, *x*_*n*_}, where *x*_*i*_ ∈ ℝ^*c×m*^ corresponds to window *i, n* is the total number of windows, and *c* is the number of epigenomic tracks. A series of four convolution modules were used to featurize each window into a vector of dimension *d* = 900 (after flattening), where each convolution module consists of a convolutional layer with ReLU activation, max pooling, and dropout. We define *Z* = {*z*_1_, …, *z*_*n*_} as the flattened output of the final convolution module where *z*_*i*_ ∈ R^*d*^ is the representation for window *x*_*i*_.

#### Bi-LSTM layer

The Bi-LSTM layers receive sequence *Z* = {*z*_1_, …, *z*_*n*_} as an input and generate a new Sequence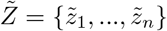, where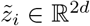. To produce the final output, every element of 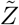 is passed through a fully connected layer yielding the output sequence *Ŷ* = {*ŷ*_1_, …, *ŷ*_*n*_}, Each *ŷ*_*i*_ ∈ ℝ^*d′*^ is a vector of dimension *d′* = 100 (or *d′* = 200 if predicting 5 kb resolution Hi-C) and corresponds to a *zig-zag* stripe in a Hi-C matrix, similar to DeepC (shown in **Fig. 1**). Epiphany uses multiple Bi-LSTM layers, with skip connections between successive layers.

#### Adversarial loss

Generative adversarial networks (GAN) consist of two networks, a generator with parameters *θ*^*𝒢*^ and a discriminator *𝒟*with parameters *θ*^*𝒟*^, that are adversarially trained in a zero-sum game [22, 35]. During training, the generator learns to fool the discriminator by synthesizing realistic samples from a given input, while the discriminator learns to distinguish real samples from synthetic samples. To train Epiphany, we employed a convex combination of pixel-wise MSE and adversarial loss. Given a dataset *D* and a trade-off parameter *λ*, Epiphany solves the following optimization problem during training:

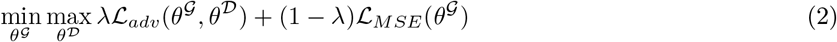

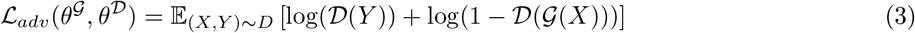

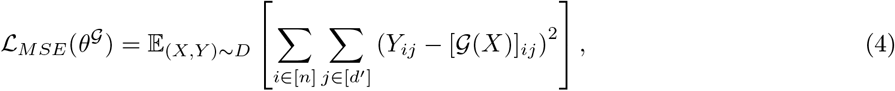

where *X* corresponds to epigenomic tracks and *Y* the corresponding Hi-C matrix.

In our framework, *𝒢* is the CNN-LSTM architecture described in the previous sections while *𝒟* is a simple four layer 2D CNN. Note that in practice, many tricks and heuristics are used in order to speed up convergence when training GANs, as described below.

#### Training

In Algorithm 1, we show the specific procedure used to approximately solve the optimization problem described above. Note that rather than setting ℒ_*D*_ to −ℒ_*G*_, we employ the target flipping heuristic outlined in [22] (Section 3.2.3) for faster convergence. The parameter updates (lines 5 and 8) are computed via the Adam optimizer [36]. We determine when to conclude training based on when ℒ_*G*_ ceases to decrease.

##### Algorithm 1

Epiphany Training

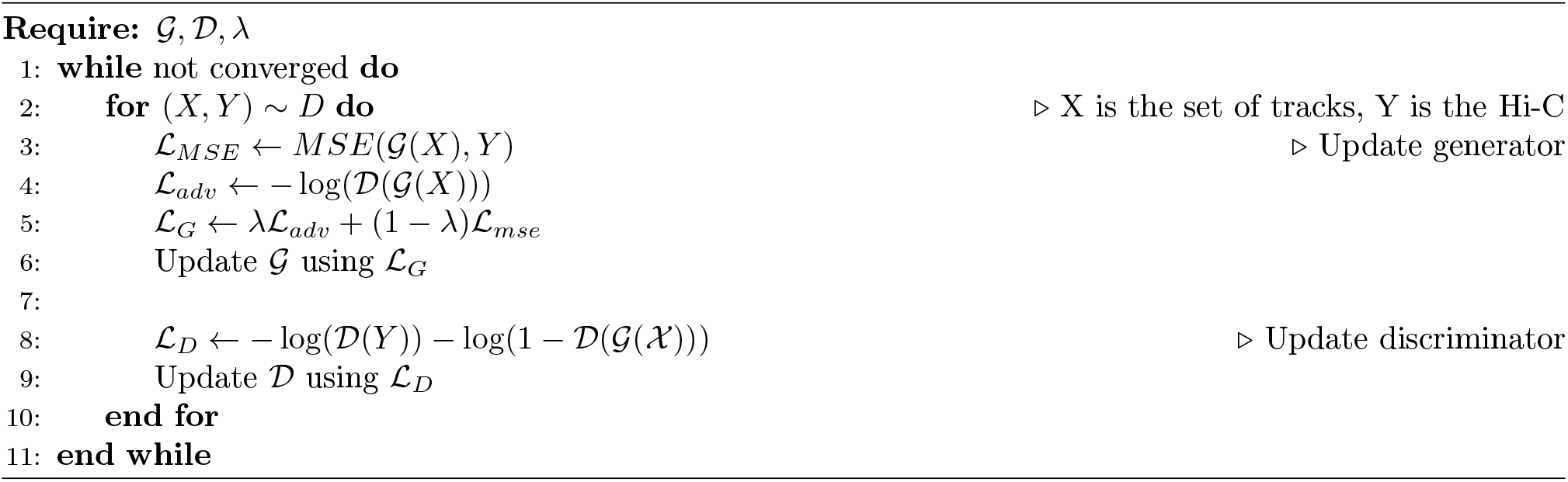

### Performance evaluation and application

#### Model performance

We evaluated the model performance using Pearson and Spearman correlation of the predicted contact map vs. ground truth, computed as a function of genomic distance from the diagonal. Predicted contact maps were saved as .hic files for downstream analysis. We visualized Hi-C matrices and epigenomic tracks using CoolBox [34]. The insulation score was calculated using the TAD-separation score from HiCExplorer [37]. Then a correlation of these scores between ground truth vs. predicted contact maps was calculated. For each 2Mb length submatrix (200 bin matrix), we calculated MSE loss and insulation score correlation between the predicted and true maps.

#### TAD boundaries and significant interactions

We identified TAD structures and significant interactions in the predicted contact maps vs. ground truth. TAD structures were identified using TopDom [23], with various window sizes of 10, 20, 30, 40, 50. Since the binary TAD boundaries would be less robust towards hyper-parameter selection and small value perturbations on the contact maps, we used insulation score for the comparisons. In this evaluation, we first ran TopDom on all ground truth contact maps with different normalization methods, and used KR normalization as the gold standard, to compare the agreement between these normalization approaches (e.g. ICE vs. KR, Z-score vs. KR). We then compared TopDom results called from predicted contact map vs. the ground truth with their corresponding normalization approach (e.g. predicted Z-score vs. ground truth Z-score), to evaluate Epiphany’s ability to predict key structures. These experiments were all run on test chromosomes 3, 11, 17.

HiC-DC+ [17] used interaction bin counts to fit a negative binomial regression with genomic distance, GC content, mappability and effective bin size based on restriction enzyme sites, providing an estimated expected read count for each interaction bins. Z-scores and observed-over-expected count ratios are then computed to evaluate the significance of the observed counts. We defined significant interactions as ground truth Z-score greater than or equal to 2. For test chromosomes 3, 11, 17 with Z-score and observed-over-expected count ratio normalization, we called significant interactions with various cut-off thresholds ranging from 0.5 to 3.5 and plotted the ROC curve.

#### Comparison with Akita

We followed provided tutorials and extracted the pre-trained Akita model from ([Akita repository](https://github.com/calico/basenji/tree/master/manuscripts/akita)). Hi-C contact maps were first balanced using ICE normalization, followed by additional steps including adaptive coarse-grain, distance adjustment, rescaling and 2D Gaussian filter suggested by Akita. Test matrices were extracted from Akita held-out test regions that overlapped with Epiphany’s test chromosomes (42 regions in total). For calculating the Pearson correlation between the predicted contact map vs. ground truth, we average-pooled Akita matrices from 2048 bp into 4096 bp, in order to keep relative consistency with our 5 kb resolution. For extracting cell-type-specific predictions, we extracted the multi-task output from Akita for H1ESc and GM12878.

#### Prediction of cell-type-specific structurea

In these experiments, Epiphany was trained on the training chromosomes in GM12878, and tested on test chromosomes chr 3, 11, 17 on H1ESc and K562. Therefore, the predictions were cross-chromosome and cross-cell-type. Feature attributions were calculated using Captum [38], with saliency score to show the gradient attribution on input regions and SHAP values to calculate the contribution of specific epigenomic peaks for predicting 3D structure. The baseline was set to zero when calculating SHAP values.

#### Feature ablation models

Feature ablation experiments were performed by re-training the model with one or several input epigenomic tracks completely masked as zero. We tested three ablation models: CTCF masked; DNaseI masked; only CTCF and H3K27ac not masked. In addition, we also re-trained Epiphany with an additional SMC3 ChIP-seq track to include cohesin occupancy information. Whole chromosome predictions were generated with trained models and compared to ground truth using Pearson and Spearman correlations as a function of genomic distance. Feature attributions were calculated as described above.

#### Biological application on mouse data

Epigenomic tracks for mouse limb bud tissue using genome assembly mm10 were downloaded from the ENCODE portal. In the CTCF deletion experiments, CTCF peaks were masked with the average value for the entire CTCF track (masked with the background). Epiphany was trained on the human cell line GM12878 and tested on mouse limb bud data (E11.5 for DNaseI and CTCF tracks, and E12.5 for H3K27ac, H3K27me3 and H3K4me3). Data were visualized using CoolBox [34].

## Data Availability

Datasets in this study are all publicly available.

### Hi-C data

Hi-C data are available from [4DN data portal](https://data.4dnucleome.org/). All human data are with hg38 assembly, and mouse with mm10.

**Table 2:**
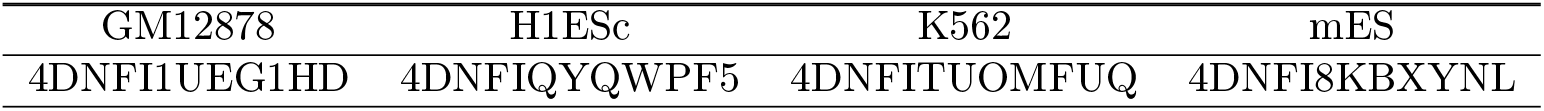
Hi-C data source

| GM12878 | H1ESc | K562 | mES |
| --- | --- | --- | --- |
| 4DNFI1UEG1HD | 4DNFIQYQWPF5 | 4DNFITUOMFUQ | 4DNFI8KBXYNL |

### Epigenomic data

Epigenomic tracks for human are available from ENCODE. Cohesin for GM12878 is available at ENCSR000DZP.

**Table 3:** Epigenomic data source

| Cell type | Dnase I | CTCF | H3K27ac | H3K27me3 | H3K4me3 |
| --- | --- | --- | --- | --- | --- |
| GM12878 | ENCSR000EMT | ENCSR000DRZ | ENCSR000DRY | ENCSR000DRX | ENCSR000AKC |
| H1ESc | ENCSR000EMU | ENCSR000AMF | ENCSR000ANP | ENCSR216OGD | ENCSR019SQX |
| K562 | ENCSR000EOT | ENCSR000DWE | ENCSR000AKP | ENCSR000AKQ | ENCSR000DWD |

### Experimental validation data

Capture Hi-C data and CTCF tracks for mES E12.5 with mm9 assembly for biological validation from Despang *et al*. [26] are publicly available at GSE78109, GSE125294.

Epigenomic tracks for mouse limb validation are available at ENCODE [27] and BioSamples [28]. All data are aligned with mm10.

**Table 4:**
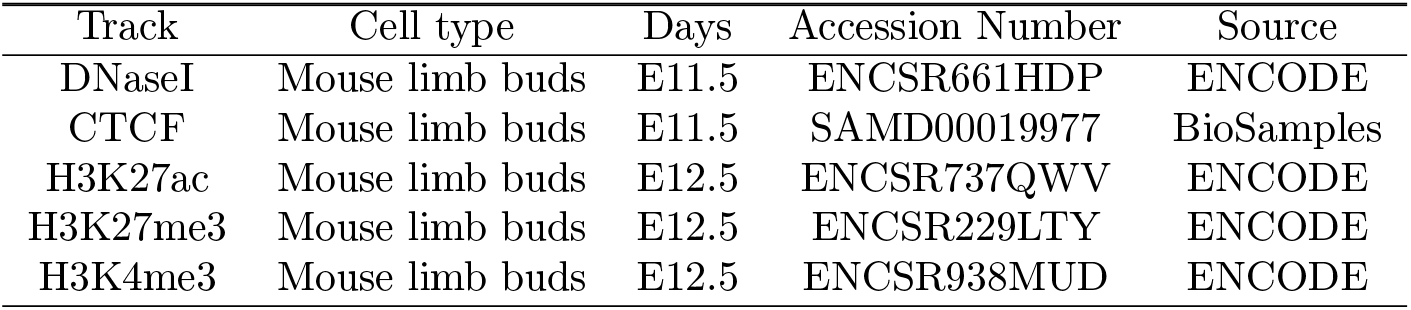
Hi-C data source

## Supplementary Information

### Model Architecture Details

In the following tables, we describe in detail Epiphany’s architecture. In each table, *n* is the total number of Hi-C stripes we seek to predict.

**Table 5:** Parameterization for 1D CNN for 5kb and 10kb Hi-C prediction

| Operation | Number of Filters | Filter Size | Stride | Activation | Output Shape |
| --- | --- | --- | --- | --- | --- |
| Input | - | - | - | - | $n \times 5 \times 14000$ |
| Convolution | 70 | $5 \times 17$ | 1 | ReLU | $n \times 70 \times 13984$ |
| Max Pool | 1 | $1 \times 4$ | 1 | - | $n \times 70 \times 3496$ |
| Dropout ($p = .1$) | - | - | - | - | $n \times 70 \times 3496$ |
| Convolution | 90 | $70 \times 7$ | 1 | ReLU | $n \times 90 \times 3490$ |
| Max Pool | 1 | $1 \times 4$ | 1 | - | $n \times 90 \times 872$ |
| Dropout ($p = .1$) | - | - | - | - | $n \times 90 \times 872$ |
| Convolution | 70 | $90 \times 5$ | 1 | ReLU | $n \times 70 \times 868$ |
| Max Pool | 1 | $1 \times 4$ | 1 | - | $n \times 70 \times 217$ |
| Dropout ($p = .1$) | - | - | - | - | $n \times 70 \times 217$ |
| Convolution | 20 | $70 \times 5$ | 1 | ReLU | $n \times 20 \times 213$ |
| Adaptive Max Pool | 1 | - | 1 | - | $n \times 20 \times 45$ |
| Dropout ($p = .1$) | - | - | - | - | $n \times 20 \times 45$ |
| Flatten | - | - | - | - | $n \times 900$ |

**Table 6:** Parameterization for Bi-LSTM for 10kb Hi-C prediction

| Operation | Hidden Layer Size | Activation | Output Shape |
| --- | --- | --- | --- |
| Bi-LSTM | 1200 | ReLU | $n \times 2400$ |
| Bi-LSTM | 1200 | ReLU | $n \times 2400$ |
| Bi-LSTM | 2400 | ReLU | $n \times 2400$ |
| Dense | - | ReLU | $n \times 900$ |
| Dense | - | None | $n \times 100$ |

**Table 7:** Parameterization for Bi-LSTM for 5kb Hi-C prediction

| Operation | Hidden Layer Size | Activation | Output Shape |
| --- | --- | --- | --- |
| Bi-LSTM | 2400 | ReLU | $n \times 4800$ |
| Bi-LSTM | 2400 | ReLU | $n \times 4800$ |
| Dense | - | ReLU | $n \times 1200$ |
| Dense | - | None | $n \times 200$ |

